# PlateletBase: A Comprehensive Knowledgebase for Platelet Research and Disease Insights

**DOI:** 10.1101/2024.12.11.627883

**Authors:** Huaichao Luo, Changchun Wu, Sisi Yu, Hanxiao Ren, Xing Yin, Ruiling Zu, Lubei Rao, Peiying zhang, Xingmei Zhang, Ruohao Wu, Ping Leng, Kaijiong Zhang, Qi Peng, Bangrong Cao, Rui Qin, Hulin Wei, Jianlin Qiao, Shanling Xu, Qun Yi, Yang Zhang, Jian Huang, Dongsheng Wang

## Abstract

Platelets are vital in many pathophysiological processes, yet there is a lack of a comprehensive resource dedicated specifically to platelet research. To fill this gap, we have developed PlateletBase, a knowledge base aimed at enhancing the understanding and study of platelets and related diseases. Our team retrieved information from various public databases, specifically extracting and analyzing RNA-seq data from 3,711 samples across 41 different conditions available on NCBI. PlateletBase offers six analytical and visualization tools, enabling users to perform gene similarity analysis, pair correlation, multi-correlation, expression ranking, clinical information association, and gene annotation for platelets. The current version of PlateletBase includes 10,278 genomic entries, 31,758 transcriptomic entries, 4,869 proteomic entries, 2,614 omics knowledge entries, 1,833 drugs, 97 platelet resources, 438 diseases/traits, and six analysis modules. Each entry has been carefully curated and supported by experimental evidence. Additionally, PlateletBase features a user-friendly interface designed for efficient querying, manipulation, browsing, visualization, and analysis of detailed platelet protein and gene information. Case study results, such as those from gray platelet syndrome and angina pectoris, demonstrate that this tool can aid in identifying diagnostic biomarkers and exploring disease mechanisms, significantly advancing research in platelet functionality and its applications. PlateletBase is accessible at http://plateletbase.clinlabomics.org.cn/.

## Introduction

Platelets, which are anucleate cellular components, are generated by megakaryocytes in the bone marrow and lungs [1]. The primary function of platelets in the body is predominantly associated with the maintenance of normal blood flow through a process known as hemostasis, which plays a crucial role in preventing excessive bleeding or hemorrhaging within blood vessels[2]. Over the past decade, numerous studies have elucidated the emerging roles of platelets beyond hemostasis and thrombosis[3]. Physiologically, extensive molecular and functional research has unveiled the multifaceted roles of platelets in preserving vascular integrity and facilitating vascular remodeling, modulating immune responses, aging and fostering tissue regeneration[4–7]. In addition to the well-established significance of platelets in cardiovascular diseases, their critical involvement in the pathophysiology of cancer, inflammatory diseases, and infections has also been extensively demonstrated[3, 8–11] (**Figure 1A**).

**Figure 1.**
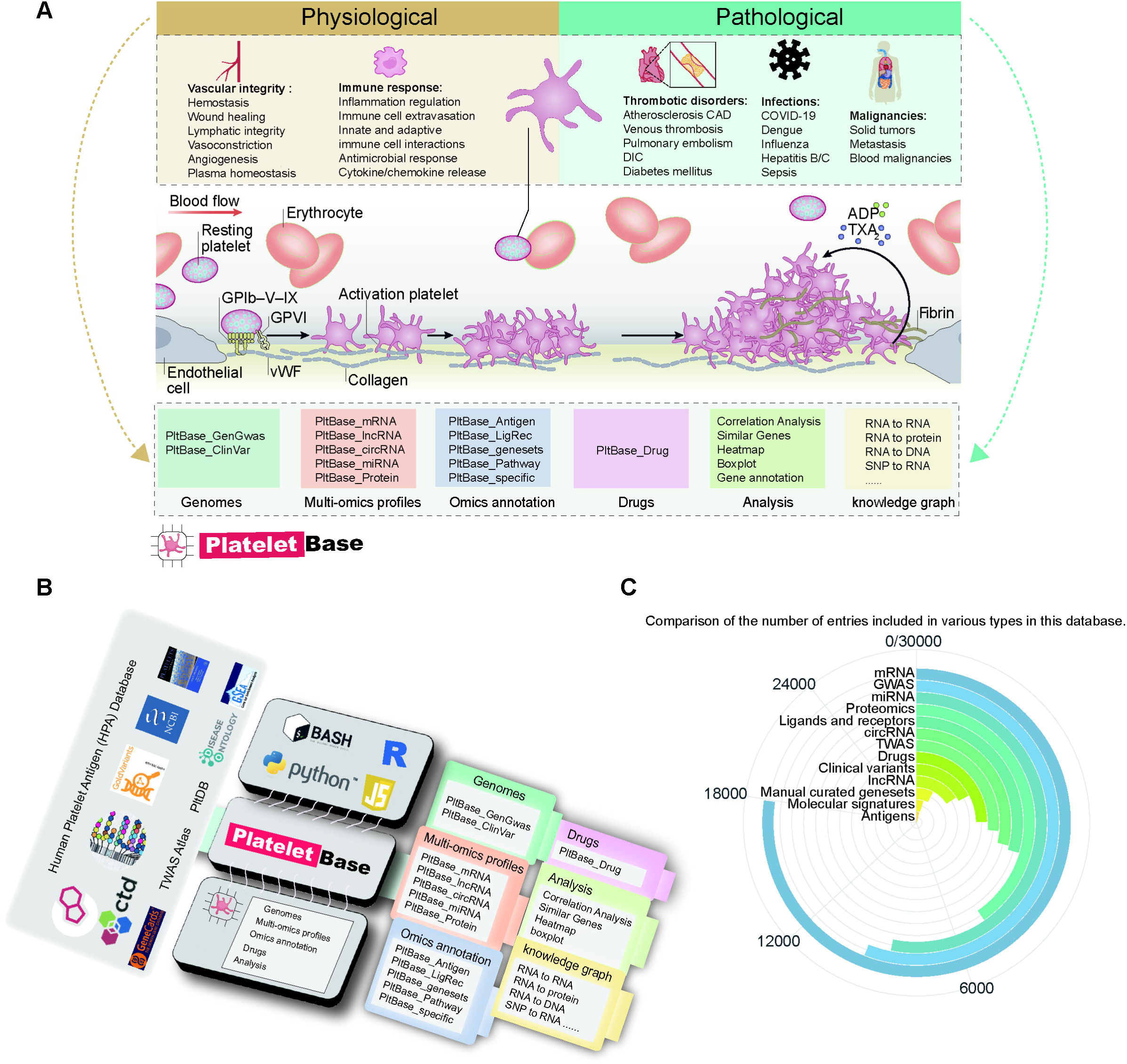
Database contents and features. **A**. The function of platelets in physiological and pathological processes. **B**. Schematic diagram of the database construction process. **C**. Entry statistics of various data types.

Despite their small size and anucleate status, platelets possess a diverse array of RNAs, including messenger RNAs, structural and catalytic RNAs, and regulatory RNAs. These RNAs play crucial roles in identifying disease biomarkers, explaining genetically or environmentally induced changes in platelet function, and facilitating the translation of mRNA into proteins, as well as transferring RNA to recipient cells to regulate various functional processes[12]. The study of mRNA and microRNA has led to the emergence of “plateletomics”[13, 14]. Research into human platelet microRNA-mRNA networks associated with age and gender has unveiled significant insights through integrated plateletomics, highlighting the intricate regulatory roles these RNAs play in platelet biology.

Nowadays, there has been a surge in research focusing on platelets-related omics. This encompasses various investigative approaches, including transcriptomics, proteomics, Genome-wide association studies (GWAS), and Transcriptome-wide association studies (TWAS), etc. Furthermore, using high-throughput sequencing and mass spectrometry technologies, researchers have undertaken significant efforts to unravel the intricate connections between genes, proteins, platelets, and diseases at various omics levels[15–20]. The primary goal of these studies is to deepen our understanding of platelets and their involvement in various diseases. By unraveling the intricacies of platelet biology, researchers aim to accelerate the development of innovative applications in diagnostics, prognostics, and therapeutics, ultimately improving patient care.

However, the retrieval, integration, and visualization of these crucial disease-gene associations, along with omics datasets, pose a formidable challenge. Valuable information is scattered across numerous scientific publications, making the process laborious and time-consuming.

Despite the existence of various resources serving different purposes, a comprehensive dedicated resource specifically for platelets is still lacking. Many of the existing resources only provide one type of omics data, limiting their scope and usefulness. Specifically, the platelet expression atlas (PEA) aims to provide comprehensive platelet expression database for diseases[21], the blood platelets-based gene expression database (PltDB) also focuses on platelet expression and offers visualization of transcriptomics profiles [22], the human platelet repository (PlateletWeb) provides a novel systems biology workbench for the analysis of platelet signaling in the functional context of integrated networks[23], Human Platelet Antigen (HPA) serves as the principal site for cataloging the current list of human platelet alloantigens. Therefore, there is an urgent need for a comprehensive and meticulously curated repository of robust disease-gene associations that encompasses a wide range of platelet-related disorders and diseases.

Here, we present PlateletBase (http://plateletbase.clinlabomics.org.cn/), a comprehensive knowledgebase specifically curated for platelet-related disorders. PlateletBase is designed for the curation, integration, visualization, and analysis of platelet and disease-related knowledge. Setting itself apart from existing databases, PlateletBase offers an extensive collection of high-quality disease-gene associations, drug-target interactions, and multi-omics datasets. This platform not only aggregates a wealth of valuable associations and interactions, meticulously curated from published literature, but also integrates a wide range of molecular profiles obtained through multi-omics data analysis. As a result, PlateletBase promises to be an invaluable resource for uncovering the molecular mechanisms driving the progression of platelet-related disorders and diseases.

## Data collection and processing

### Knowledge curation and integration

Platelet-related genomic alterations are collected from GWAS Catalog[24], TWAS Atlas[25], ISTH Gold Variants project[26]. To obtain comprehensive data on platelet antigens and antigen genes, the Human Platelet Antigen (HPA) Database and HPA Gene Database were used as sources. The information from these two databases was integrated into a single statistical platelet antigen dataset, providing systematic coding. In order to obtain pathways relevant to platelets, the latest version of the MSigDB database, msigdb_v2023.2, containing tens of thousands of annotated gene sets for use with GSEA software, was downloaded. Keywords such as ‘Platelet,’ ‘Thrombocyte,’ and ‘coagulation’ were utilized in the search. To assemble a collection of platelet-related ligand-receptor pairs, we initially employed the liana R package to extract and amalgamate ligand-receptor pairs from 77 databases such as “CellTalkDB”, “CellChatDB,” “CellPhoneDB,” and “CellCall,” supported by experimental evidence and manual curation[27–29].

To obtain data on the association between platelet proteomics and diseases, extensive literature searches were conducted using the keywords “Proteomics” and “platelet.” Eventually, 25 relevant studies were compiled, covering 21 different diseases with a total sample size of 1,185 individuals. Referencing the book “Platelet”, three classic articles on platelet proteomics were identified, focusing on resting-state platelets, activated platelets, and platelet membrane proteins, which serve as reference data. The integration of these reference data with the compiled literature yielded a total of 4,869 different types of proteins identified in platelet proteomics. For better understanding, proteins were numbered based on their expression levels in resting state, with annotations indicating their expression levels under different conditions[30–32].

To obtain data on the association between platelets, drugs, and genes, we conducted searches in three authoritative databases for drug-gene associations using multiple platelet-related keywords: CTD [33], ChEMBL[34], and PharmGKB[35]. Through manual curation, we compiled 1833 entries of platelet-related drug information, involving 80 genes, 148 diseases, and 225 articles.

To coordinate disease nomenclature and definitions, terminology or identifiers from the Disease Ontology were retrieved and mapped to the corresponding diseases[36]. In order to provide consistent names for all collected genes, gene names were unified with the help of the gene symbol-alias conversion table from the HGNC database (2021.4.23 version) [37].

### Transcriptome Data collection and Raw data processing

The platelet transcriptome refers to the set of all RNA molecules within platelets. Although platelets are anucleate cells, they possess a rich array of RNA, including messenger RNA (mRNA), microRNA(miRNA), long non-coding RNA (lncRNA), circular RNA (circRNA), etc. To obtain data on the association between the platelet transcriptome and diseases, we searched public databases and relevant literature for platelet-related RNA-seq raw data. All collected data were processed through a consistent bioinformatics pipeline. First, the quality of the raw RNA-seq Fastq data was assessed using FastQC (v0.11.9) and MultiQC (v1.13)[38]. Based on these results, low-quality reads and adapter sequences were filtered out using Fastp (v0.22.0) to obtain clean reads. Next, STAR (v2.7.10b) was used to align the clean reads to the human reference genome (GRCh38), and quantification was performed using HTSeq (v2.0.2) to convert the aligned reads into count values for each gene. The expression values were then normalized to TPM, and Ensembl Gene IDs were converted to Gene Symbols, retaining the one with the highest mean expression if multiple Gene IDs corresponded to the same Gene Symbol. This process yielded processed mRNA and lncRNA data.For circRNA detection, the clean reads were additionally aligned using BWA (v0.7.17) and detected using CIRI (v2.0.6)[39]. Expression values were normalized to RPM and similarly converted to Gene Symbols to obtain the final circRNA data.

For mRNA, to ensure gene generality and comparability, genes missing in more than 25% of patients were deleted, while those missing in less than 25% were filled with 0. After applying log2(TPM+1) transformation, the ComBat function from the SVA R package was used to remove batch effects, considering different project sources as batch factors. The curated list of all Hugo genes was used as the standard gene names, and genes were numbered based on their expression levels from high to low in 802 normal samples, resulting in a total of 17,812 genes.

To identify differential mRNA between each physiological or pathological state and the normal group, the Wilcoxon Rank Sum Test was used for each disease group compared to the normal group, with thresholds of a p-value of 0.05 and a logFC of 0.5, identifying 8,858 differentially expressed mRNA across various disease states. To calculate characteristic mRNA for each physiological state, we considered any given condition as the candidate state (case group) and all other groups as the control group, and we calculated specific mRNA for different groups. Using the Wilcoxon Rank Sum Test with thresholds of a p-value of 0.05 and a logFC of 0.5, we identified the specific genes. For other RNAs, please refer to the corresponding sections of the website for analysis methods.

For miRNAs, we extracted the batch-corrected miRNA dataset from PltDB[22]. By ranking the miRNAs based on their expression levels in the control group, we defined a total of 8,468 miRNAs. This dataset involves 167 samples and covers 6 different diseases. To identify differential miRNA between each physiological or pathological state and the normal group, the Wilcoxon Rank Sum Test was used for each disease group compared to the normal group, with thresholds of a p-value of 0.05 and a log2 Fold change (logFC) of 0.5.

Ultimately, our platelet transcriptome dataset covers 41 pathophysiological conditions with a total of 4408 samples. The final dataset comprises 17,926 types of mRNA, 756 types of lncRNA, 2493 types of circRNA, and 8468 types of miRNA, all expressed across various physiological and pathological states.

### Database construction and web interface implementation

PlateletBase is built using Spring Boot (https://docs.spring.io/spring-boot/index.html), Mybatis (https://mybatis.net.cn/), and MySQL (http://www.mysql.org). Its web interface is developed with CSS3, Axios, Element-UI (version 2.15.8), Vue-Router (version 3.0.6), and Vue (version 2.x). Comprehensive statistical visualization of the data is achieved through HTML widgets, R, and Python, primarily utilizing ggplot2 (version 3.5.1) and Plotly (version 5.19.0) (**Figure 1B**).

### Knowledge graph construction

To better illustrate the correlations among various omics of platelets, this analysis module incorporates the established relationships between RNA and proteins, DNA, as well as RNA identified in experiments or literature from the RNAInter v4.0 dataset[40]. Additionally, we integrate eQTL data (GTEx_Analysis_v10_eQTL) to establish connections between genomic information, gene expression, and tissue localization[41]. Furthermore, we have consolidated the differential expression tables of mRNA and the proteome according to gene names to effectively demonstrate the relationship between mRNA expression and protein expression within platelets.

### Analysis modules construction

The analysis section, based on Python, employed various statistical methods such as Pearson, Spearman, and Kendall correlation analyses, as well as Wilcoxon rank-sum test and Fisher’s exact test. It implemented a range of analytical tools and visualized the results using the Matplotlib package to create correlation plots, heatmaps, boxplots, and bar charts.

## Database content and usage

PlateletBase encompasses approximately 9381 GWAS entries, 2115 TWAS entries, 897 clinical variations, 17,926 mRNA entries, 756 lncRNA entries, 2493 circRNA entries, 8468 microRNA entries, 4869 proteins, 40 platelet antigens, 65 platelet-related pathways, 2614 ligand-receptor pairs, 1833 drugs, and 97 platelet resources (**Figure 1C**). These entries have been meticulously curated, ensuring that each one is supported by experimental evidence. PlateletBase offers a user-friendly interface tailored for efficient querying, manipulation, browsing, visualization, and analysis of intricate details regarding platelet proteins and genes. This resource promises to significantly advance research endeavors in platelet functionality and applications.

### Platelet-related genomic alterations

Platelet-related traits have a high genetic heritability, with up to 84% of the variance in platelet count and 75% of the variance in mean platelet volume (MPV) attributed to genetic factors[42]. Early extensive research has studied the relationship between various platelet phenotypes and genetics. The Manhattan plot in **Figure 2A** shows that these platelet phenotypes do not seem to prefer chromosomal distribution. PlateletBase has integrated 112 genetically-related platelet phenotypes, such as platelet aggregation, platelet count, mean platelet volume, etc., among which the most important phenotypes are platelet count and MPV. The three SNP loci most frequently linked to phenotypes are rs1354034 (28 phenotypes), rs385893 (17 phenotypes), and rs11082304 (13 phenotypes) (**Figure 2A**). The top three most frequently occurring pairs of phenotypes and SNP loci are Platelet count - rs1354034, Platelet count - rs11082304 and Platelet count - rs11082304 (**Figure 2B**). Studies have shown that the rs1354034 mutation can affect platelet function and be associated with ischemic stroke subtypes[42, 43]. The minor allele mutation of rs11082304 is associated with platelet count[44].

**Figure 2.** Statistics derived from platelets associated genomics datasets. **A**. Manhattan plot showing the relationship between chromosome position and -log10(p value) of all platelet-related GWAS loci, with the top 3 most frequent (Phenotype or disease involved)loci annotated in orange, and SNPs with -log10(p value) > 300 labeled. **B**. Bar graph showing the distribution of frequencies of platelet-related GWAS loci. **C**. Frequency of clinical mutations associated with platelets. **D**. The ring diagram shows the disease composition corresponding to the top 4 clinical gene mutations. **E**. Bar chart showing the distribution of relationships between platelet-related phonotypes and top 5 tissues gene expression from TWAS data.

Clinically relevant variations may significantly impact a patient’s life. GoldVariants, developed and tested by 30 expert centers, play a crucial role in identifying and validating Clinically Relevant Variations (ClinVars) [26]. The ClinVars in this database are primarily derived from the variations identified in GoldVariants. Analyses indicate that these clinical mutations predominantly arise from important platelet-related genes, such as ITGA2B, ITGB3, and GP1BA, which are critical for platelet function and associated with various bleeding disorders (**Figure 2C**).

The distribution of mutations for the four most frequently occurring clinically relevant genes is as follows: GP1BA mutations lead to Bernard-Soulier syndrome (13.8%), macrothrombocytopenia (2.3%), and mild macrothrombocytopenia (18.4%); ITGA2B and ITGB3 mutations are associated with Glanzmann thrombasthenia (96.7% and 97.5% respectively) and platelet-type bleeding disorder (approximately 3% each); VWF mutations cause von Willebrand disease (100%). These mutations affect platelet function, resulting in varying degrees of bleeding tendency (**Figure 2D**).

### Platelet transcriptome changes associated with diseases

While GWAS have uncovered numerous common genetic variants linked to complex traits, the precise causal variants and genes at these loci frequently remain elusive, with a handful of exceptions. TWAS integrate GWAS and expression quantitative trait loci (eQTL) data to forecast gene expression levels for GWAS samples and then assess for associations between the predicted expression and traits[45]. We can see some tissues are burdened with increased levels of risk genes for a given platelet traits. Eventually, 7 relevant studies were compiled. The two most frequently studied phenotypes were “mean platelet volume” and “platelet count”. Across 52 different tissues, aside from whole blood, the most frequently occurring tissues were “Testis,” “Thyroid,” and “Adipose”. Further analysis revealed that the genes related to mean platelet volume (MPV) are all expressed in the blood, while the genes associated with platelet count are expressed in various tissues (**Figure 2E**).

Research indicates that platelet mRNA can be translated into proteins or transferred to recipient cells, thereby regulating functional processes. This database extracted 20 sets of original data, 3711 samples, and 41 different conditions from NCBI (**Figure 3A**).

**Figure 3.** Statistics derived from curated platelets transcriptomics and proteomics data collection. **A**. Sankey diagram showing the disease distribution of the RNA data sets. **B**. Heatmap of feature gene expression in platelets for several major disease types. **C**. Bar chart of enriched genes that are highly expressed in platelets from healthy individuals. **D**. Sankey diagram showing the disease distribution of the proteomics expression data

In order to better demonstrate the disease-specific expression of platelet-related RNAs, we extracted the 3 genes with the greatest differential expression for each major disease category, and displayed them in a heatmap. The inherent similarities and differences between diseases can be well reflected by the differential platelet expression profiles. The tumor diseases form a distinct cluster, while the viral infections and immune system diseases form a similar branch (**Figure 3B**). We enriched the top 150 genes highly expressed in the healthy group through Metascape, and found normal cell functions such as gas transport and positive regulation of cell adhesion[46] (**Figure 3C**).

### Platelet proteome changes associated with diseases

The composition, localization, and activity of proteins are essential for platelet function and regulation. The current advancements in mass spectrometry-based proteomics offer significant potential to detect and quantify thousands of proteins from minimal sample quantities, uncover various post-translational modifications, and track platelet activity in response to drug treatments. This database extracted data from 25 publications covering 21 different diseases, identifying 1,345 differentially expressed proteins. Sankey diagram analysis shows that the majority of highly expressed proteins in cancers and cardiovascular diseases are primarily derived from activated platelets (**Figure 3D**).

### Platelet-related antigens

Alloantibodies against human platelet antigens (HPA) are implicated in several immune-mediated platelet disorders, such as fetal and neonatal alloimmune thrombocytopenia and platelet transfusion refractoriness. The detection and identification of these HPA alloantibodies are critical for accurate diagnosis and the implementation of appropriate patient care strategies. In PlateletBase, 40 distinct platelet antigens were compiled, with 32 located on chromosome 17, as referenced in 54 publications.

### Platelet-related molecular signatures and manual curated gene sets

MSigDB is a comprehensive resource that houses a collection of gene sets representing specific biological pathways, molecular processes, and cellular functions. These gene sets are curated from diverse sources, including experimental studies, computational predictions, and literature annotations. The database facilitates the systematic exploration of gene expression data by allowing researchers to analyze and interpret their results in the context of known biological pathways and signatures. Ultimately, 65 platelet related molecular signatures were compiled, involving 2,056 genes, which are displayed by word cloud (**Figure 4A**).

**Figure 4.** Statistics derived from omics knowledge and drugs associated with platelets. **A**. The word cloud displays all platelet-related molecular signaling pathways in the database, with the size and color of the pathway proportional to the number of genes in the pathway. **B**. Bar chart showing the top 10 most frequent Ligand gene symbols. **C**. Bar chart showing the top 10 most frequent Receptor gene symbols. **D**. Bar chart showing the top 10 most frequent platelet associated drug gene symbols. **E**. Venn diagram showing gene overlap.

To provide more molecular biology information for platelet research, we compiled 15 datasets related to platelet production, adhesion, reticulated platelets, and other relevant areas, involving 194 genes or proteins. Additionally, using a method similar to the MSigDB, we sorted 150 (or fewer if the criteria were not met) highly or lowly expressed genes by logFC, ensuring p-values were less than 0.05 in platelet mRNA transcriptome data. Collectively, we compiled 82 gene sets associated with 41 disease states, based on differentially expressed genes compared to normal platelets. Furthermore, we compiled disease-specific gene sets by comparing various diseases and other conditions, resulting in 84 gene sets across 42 conditions, including normal states.

### Platelet-related ligands and receptors

Platelet receptors play crucial intercellular communication roles by serving as both ligands and receptors, facilitating interactions between platelets and other cells or extracellular molecules. These receptors mediate signaling pathways that regulate platelet activation, adhesion, and aggregation in response to external stimuli. By recognizing and binding to specific ligands, such as adhesive proteins or agonists, platelet receptors initiate cellular responses that contribute to hemostasis, thrombosis, and other physiological processes. Through these intercellular communication mechanisms, platelet receptors coordinate complex cellular interactions within the vascular system. In PlateletBase, we compiled a comprehensive set of 2614 ligand-receptor pairs, encompassing 770 receptors and 808 ligands. Among these, 77 pairs had been previously reported, while 1528 receptor gene expressions in platelets and 1009 ligand gene expressions in platelets were novel discoveries. Top gene are displayed by bar plots (**Figure 4B-C**).

### Platelet-related drug-target interactions

Understanding medications for platelet-related diseases is crucial for effective management. These drugs, such as antiplatelet agents and anticoagulants, play a vital role in preventing thrombosis and managing cardiovascular conditions. Genetic factors can influence how individuals respond to these medications, highlighting the importance of personalized treatment approaches. However, emphasizing the significance of platelet-related medications underscores their direct impact on patient care and clinical outcomes. Through manual curation, we compiled 1833 entries of platelet-related drug information, involving 80 genes, 148 diseases, and 225 publications. CD36 is highest frequency gene, which also called platelet glycoprotein IV, acts as both a signaling receptor and a transporter for long-chain fatty acids[47] (**Figure 4D**). We further applied Venn diagram analysis to display the gene overlap among the different omics data. We found that the gene sets for ‘Platelet-related ligands and receptors’ and ‘Platelet-related molecular signatures’ shared the largest number of common genes (**Figure 4E**).

### Personalized analysis module for platelet omics

This database has detailed help documentation. It provides a simple introduction to the navigation bar, where the standout feature is the analysis module, which facilitates customized analysis by researchers to obtain more information. Compared to previous platelet-related databases, our database is the first platelet multi-omics database, and the sample size of various omics data exceeds that of previous databases, with more disease types covered (**table 1**).Based on the transcriptome data in the database and backend programs, this database provides six analysis and visualization methods: similar gene analysis by correlation allows users to select a correlation method, disease types, gene of interest, and result proportion to see genes most correlated with the input gene; pair correlation enables users to analyze the correlation between two genes in selected diseases, with results shown in a scatter plot; Multi-Correlation lets users examine correlations among multiple genes, with results in a correlation heatmap; expression rank in all conditions allows users to view a gene’s expression across diseases and healthy samples in a box plot; and clinical analysis enables users to explore a gene’s expression under different clinical conditions within selected diseases, presented in a box plot. Users can choose the RNA type (mRNA, lncRNA, circRNA, miRNA) and the appropriate analysis method to get the corresponding results.

The users also can conduct enrichment analysis. This function is designed to analyze whether specific genes or molecules are significantly enriched in certain biological contexts, helping users quickly identify potential key biological mechanisms or pathways. Users can input a gene list of interest and select relevant gene sets from the database based on their research needs (with the option to select all). Enrichment analysis is then performed using Fisher’s exact test, and the results are presented in a tabular format, including the following information: gene set name, the number of genes in the gene set, the number of overlapping genes, the proportion of overlapping genes, p-value, and adjusted p-value. Additionally, a bar chart visualizing the top 10 most significantly enriched gene sets is generated for better interpretation.

### Integrate multi-omics information through knowledge graph

Considering the pertinent isolation of multi-omics data, it is imperative to establish connections among various data types. This section is organized into three principal modules. The first module employs RNA as the initial point for interactions. Users may select from three distinct interaction modalities: RNA-RNA, RNA-DNA, and RNA-Protein. Upon entering a gene name, users can obtain both graphical and tabular representations illustrating the gene’s involvement in network interactions across different forms. The second module is centered on SNP loci, aiming to elucidate the functional associations between SNPs correlated with platelet phenotypes and gene expression within specific tissues. Users can input a gene, SNP locus, or tissue name to generate corresponding network relationship diagrams and tables. The final module integrates tables of platelet RNA and platelet proteins, utilizing gene symbols as linking entities. This facilitates users in exploring the associations between the same gene in both RNA and protein forms and their relevance to various diseases.

### Case study 1: Gray platelet syndrome

Previous studies have indicated that gray platelet syndrome (GPS) is a rare platelet disorder characterized by abnormalities in α-granules and mutations in NBEAL2, featuring macrothrombocytopenia and a specific deficiency of α-granules and their associated contents.[48]. A user seeking to understand gray platelet syndrome (GPS) can first navigate to the genomics section and click on the clinical variant subsection, selecting GPS. This will reveal 16 distinct mutations associated with the pathogenesis of GPS. Consistent with prior research, all identified mutations involve the NBEAL2 gene. By accessing the hyperlink for this gene, users can explore more in-depth studies. GWAS data indicate that the SNP loci of the NBEAL2 gene are correlated with mean platelet volume characteristics, aligning with the phenotypic features of GPS (**Figure 5A**). In the omic knowledge section, NBEAL2 is associated with α-granules, appearing within several platelet-related gene sets (**Figure 5B**).

**Figure 5.** Case report of Gray platelet syndrome. **A.** Basic information about the main gene NBEAL2 related to Gray platelet syndrome and GWAS-related locus information. **B.** Molecular pathway information associated with NBEAL2. **C.** List of drugs related to NBEAL2. **D.** Protein network diagram related to NBEAL2.

In the pharmacological section, multiple therapeutic agents targeting the NBEAL2 gene for the treatment of GPS are listed (**Figure 5C**). Furthermore, transcriptomic and proteomic analyses and Knowledge graph related to NBEAL2 (**Figure 5D**) have revealed additional relationships between NBEAL2 and various diseases and proteins, potentially offering new avenues for mechanistic studies or repurposing existing medications.

### Case study 2: Angina pectoris

Angina significantly contributes to global morbidity, with its assessment being variable and influenced by multiple factors. Cardiac troponins have replaced other blood biomarkers for diagnosing myocardial injury [49]. However, unstable angina is less frequently diagnosed with high-sensitivity cardiac troponin (hs-cTn) assays[50]. This study aims to identify platelet-based diagnostic biomarkers for angina using this database. Initially, we accessed the transcriptomics section to identify differentially expressed genes associated with angina compared to healthy individuals, as well as specific differential genes compared to non-angina patients, sorting them by log2 fold change (log2FC). We selected the gene with the highest log2FC, identifying CAPN10 as a candidate biomarker.

We examined the basic information for this gene through its hyperlink (Figure 6A) and confirmed the validity of the differential expression using the built-in plotting tool (Figure 6B). To consider CAPN10 as a biomarker, we needed to identify potential differential diagnosis factors; therefore, we ranked diseases based on CAPN10 expression. We found that patients with unstable angina exhibited the highest platelet expression of CAPN10 among over 40 diseases, necessitating differentiation from only a few conditions (Figure 6C).

**Figure 6.** Case report of Angina Pectoris. **A.** Basic information about the differentially expressed gene CAPN10 related to angina pectoris. **B.** Boxplot comparing expression levels in angina pectoris (left) versus healthy individuals (right). **C.** CAPN10 expression rank across all conditions. **D.** The top 100 genes similar to CAPN10, enriched through Enrichment Analysis and Plot function in PlateletBase, related to platelet molecular signaling pathways.

To further investigate the potential function of this gene in platelets, we utilized the similar genes module to obtain the top 100 similar genes and employed the enrichment module, revealing that CAPN10-related genes are primarily enriched in gene sets associated with platelet activation (Figure 6D). Therefore, this tool can be utilized for the exploration of diagnostic biomarkers.

## Discussion and perspectives

In this study, we present PlateletBase, a comprehensive knowledge base that provides a high-quality collection of data on human platelets, associated diseases, genes, drugs, and omics profiles. Based on integration from published literature and related databases, the current version of PlateletBase contains 10,278 genomics entries, 31,758 transcriptomics entries, 4,869 proteomics entries, 2,614 omics knowledge entries, 1,833 drugs, 97 platelet resources, 438 diseases/traits and 6 analysis modules. Future directions include: (i) frequent curation and expansion of all data sections; (ii) integration and analysis of additional multi-omics datasets for platelets; and (iii) improvement of the web interface and development of tools to facilitate multi-omics data mining and visualization. It is important to note that many aspects of this study are based on data mining, which necessitates further validation to corroborate the findings. We call on the global scientific community to collaborate in building PlateletBase into a valuable resource covering more comprehensive associations, interactions, and omics data, in order to further advance high-quality, curated knowledge for platelet and disease research.

## Data availability

PlateletBase is freely available online at http://plateletbase.clinlabomics.org.cn/ and does not require user to register.

## CRediT author statement

Huaichao Luo: Methodology, Data curation, Software, Visualization, Writing – original draft, Funding acquisition. Changchun Wu: Software, Supervision. Sisi Yu: Resources. Hanxiao Ren: Resources. Xing Yin: Resources. Ruiling Zu: Resources, Funding acquisition. Lubei Rao: Resources, Funding acquisition. Peiying zhang: Resources. Xingmei Zhang: Resources. Ruohao Wu: Resources. Ping Leng: Resources, Supervision. Kaijiong Zhang: Resources, Supervision. Qi Peng: Resources, Supervision. Bangrong Cao: Conceptualization, Supervision. Rui Qin: Software, Supervision. Hulin Wei: Methodology, Data curation, Software, Visualization. Jianlin Qiao: Conceptualization, Supervision. Shanling Xu: Conceptualization, Supervision, Funding acquisition. Qun Yi: Conceptualization, Supervision, Funding acquisition. Yang Zhang: Resources, Supervision. Jian Huang: Conceptualization, Supervision. Dongsheng Wang: Writing – review & editing, Funding acquisition. All authors have read and approved the final manuscript.

## Acknowledgments

We are grateful to all the original authors and contributors of the data involved in this database. Due to the length constraints of this article, we are unable to cite them individually, but we would like to express our sincere appreciation for their invaluable contributions. We apologize for not being able to acknowledge them one by one, and we thank them once again. This work was supported by grants from the Natural Science Foundation of Sichuan Province (Grant No. 2024NSFSC0767, 24NSFSC1880, 2024NSFSC1556), National Key Research and Development Program of China (Grant NO. 2023YFC2507200), Sichuan Provincial Health Commission Science and technology projects (Grant NO.23LCYJ041), and Noncommunicable Chronic Diseases-National Science and Technology Major Project (Grant NO.2023ZD0506603).

